# Functional Brain Network Estimation with Time Series Self-scrubbing

**DOI:** 10.1101/191262

**Authors:** Weikai Li, Lishan Qiao, Zhengxia Wang, Dinggang Shen

**Affiliations:** School of Mathematics Science, Liaocheng University, Liaocheng 252000, China and with College of Information Science and Engineering, Chongqing Jiaotong University, Chongqing 400074, China.; School of Mathematics Science, Liaocheng University, Liaocheng 252000, China.; College of Information Science and Engineering, Chongqing Jiaotong University, Chongqing 400074, China.; Department of Radiology and BRIC, University of North Carolina at Chapel Hill, NC 27599, USA and with Department of Brain and Cognitive Engineering, Korea University, Seoul 02841, Republic of Korea..

**Keywords:** Functional Brain Network, Resting-state Functional Magnetic Resonance Imaging (fMRI), Mild Cognitive Impairment (MCI)

## Abstract

Functional brain network (FBN) has been becoming an increasingly important measurement for exploring the cerebral working mechanism and mining informative biomarkers for assisting diagnosis of some neurodegenerative disorders. Despite its potential performance in discovering the valuable patterns hidden in the brains, the estimated FBNs are often heavily influenced by the quality of the observed data (e.g., BOLD signal series). In practice, a preprocessing pipeline is usually employed for improving the data quality prior to the FBN estimation; but, even so, some data points in the time series are still not clean enough, possibly including original artifacts (e.g., micro head motion), non-resting functional disturbing (e.g., mind-wandering), and new “noises” caused by the preprocessing pipeline per se. Therefore, not all data points in the time series can contribute to the subsequent FBN estimation. To address this issue, in this paper, we propose a novel FBN estimation method by introducing a latent variable as an indicator of the data quality, and develop an alternating optimization algorithm for scrubbing the data and estimating FBN simultaneously in a single framework. As a result, we can obtain more accurate FBNs with the self-scrubbing data. To illustrate the effectiveness of the proposed method, we conduct experiments on two publicly available datasets to identify mild cognitive impairment (MCI) patients from normal control (NC) subjects based on the estimated FBNs. Experimental results show that the proposed FBN modelling method can achieve higher classification accuracy, significantly outperforming the baseline methods.

## I. INTRODUCTION

FUNCTIONAL brain network (FBN) provides an increasingly important way to explore the brain integration mechanism [1], discover potential brain activities [2], and mine sensitive biomarkers for neural disease diagnosis [3,4] (such as autism spectrum disorder [5-7], Alzheimer’s disease [8,9] and Parkinson’s disease [10]). All these rely heavily on the quality of the final FBNs, and thus it becomes essential to estimate an accurate FBN [11].

Currently, researchers have proposed many different methods towards better FBN estimation, and most of them can be explained under a regularized framework that requires an accurate FBN estimation model to *not only* fit the data, but also effectively encode the priors of brain organization [12]. In practice, the commonly-used priors include sparsity [13-16], group-sparsity [17,18], low-rank [19], and modularity [12], which can be transformed into their corresponding regularization terms in the FBN estimation models and can often improve the performance of the obtained FBNs.

Besides the regularizers, the data-fitting terms (see Eq. (5) and related comments for details) also have a high influence on FBN estimation. However, the artifacts or noises involved in the observed data (or time series) usually lead to a poor fitting result. Therefore, in practice, a preprocessing pipeline, including motion correction, spatial smoothing and temporal filtering, is generally employed to improve the quality of the data before FBN estimation [20]. Even so, it is hard to eliminate all the artifacts/noises in the data due to the weak fMRI signals and complex disturbing sources. Moreover, some preprocessing steps (e.g., spatial normalization) in the pipeline may also cause new “dirty” points in the time series.

As a recently developed preprocessing step, the scrubbing operation has been investigated to further clean the data by removing some potentially “dirty” time points from the fMRI series [21]. Despite its seeming appeal, there are also some debates on such a scheme [21-25]. Moreover, this scheme mainly focuses on specific physical artifacts (e.g., micro head motion) that can only cover a limited range of problematic time points. In fact, a large number of factors may introduce “dirty” data points into the fMRI series. For example, the “resting-state” fMRI data tend to involve many different functional processes, e.g., mind-wandering [26], thus resulting in non-resting-state time points in the fMRI data that cannot be detected by the traditional scrubbing technique. More notably, the scrubbing operation is independent of the FBN estimation model used, and thus cannot guarantee that the preserved data points can necessarily benefit for the subsequent FBN estimation, while the removed data points are not necessarily helpful. In addition, it is hard for the common scrubbing method to control the length of the remaining time series [21], and thus the final series sometimes cannot contain enough time points (samples) for estimating a reliable FBN [12].

To address these issues, in this paper, we propose a novel FBN estimation strategy by introducing a latent indicator variable into the FBN optimization model. The latent variable indicates the quality or state of each time point in the fMRI series, and can be learned automatically with the FBN together from the data. Based on the latent variable model [27,28], we then develop an alternating optimization algorithm to simultaneously estimate the indicator variable and FBN in a single framework. Consequently, our proposed method will be able to automatically identify and remove the “dirty” time points from the fMRI series; that is, it can make a self-scrubbing operation at the FBN estimation procedure. In the following, the contributions and the main advantages of our proposed framework are summarized.

1) Different from traditional methods that often conduct the data scrubbing and FBN estimation in two separate sequential steps, our proposed framework combines data scrubbing and FBN estimation into a single model. By joint optimization, we can obtain FBNs with potentially higher accuracy and efficiency.

2) Technically, we introduce a latent variable into the FBN estimation model as an indicator of the data quality, and design an alternating approach to estimate the optimal indicator, by which we can scrub the time points that are potentially not helpful for FBN estimation. Moreover, compared with the traditional scrubbing operation that often removes too many data points, our proposed framework can control the size of scrubbed data by a hyper-parameter.

3) Finally, it is worth emphasizing that our proposed framework is not competitive to the original scrubbing strategy, since it can also work on the data that have been already scrubbed by the traditional methods. Therefore, the proposed method is more flexible and adaptive for the FBN estimation than the traditional scrubbing scheme.

To verify the effectiveness of the proposed method, we apply it to estimate FBNs based on the resting-state functional magnetic resonance imaging (R-fMRI), and then identify mild cognitive impairment (MCI) patients from normal control (NC) subjects via the estimated FBNs. Experiments are conducted on two publicly available datasets, and the experimental results illustrate that our proposed method works well on both scrubbed and non-scrubbed R-fMRI data. For facilitating efforts to replicate our results, we have also shared both pre-processed data and codes in https://github.com/Cavin-Lee/self-scrubbing/.

The rest of this paper is organized as follows. In Section <u>II</u>, we review the main-stream FBN estimation methods. In Section <u>III</u>, we propose a novel FBN estimation framework, including motivation, modeling, and algorithm. In Section <u>IV</u>, we conduct experiments on a simulated data and two real-world datasets. In Section <u>V</u>, we conclude the paper with a brief discussion.

## II. RELATED WORKS

In this paper, we mainly focus on the correlation-based methods, which are currently the most popular ways of FBN estimation and have been empirically demonstrated to be more sensitive than the complex or higher-order methods [11]. In the following, we review several representative correlation-based methods that, in fact, provides a platform for developing our model.

### A. Pearson’s Correlation

According to a recent review [1], Pearson’s Correlation (PC) is the simplest method for estimating FBNs. We suppose that each brain has been parcellated into *N* regions of interest (ROIs) based on a certain atlas, and the fMRI time series associated with the *i*th ROI is represented by **x**_*i*_ ϵ *R*^*T*^, *i* = 1, …, *N*, where *T* is the number of time points in each series. Then, the edge weights of the FBN based on PC can be calculated as follows:

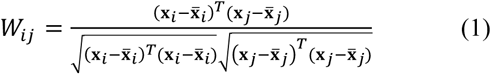

By defining a new 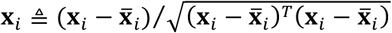, a centralized and normalized counterpart of the original X_*i*_, we can simplify the PC as *W*_*ij*_ = **x**_*i*_^*T*^**x**_*j*_, which corresponds to the solution of the following optimization problem:

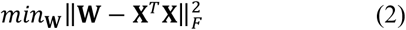

Here, **X** = [**X**_1_, **X**_2_, …, **X**_*N*_] ϵ *R*^*T×N*^ is the data matrix, **W** is the edge weight matrix, and ‖∥·‖∥_*F*_ denotes the F-norm of a matrix. Since the BOLD signals commonly contain noises, the original PC tends to result in a FBN with dense connections [29]. In practice, a threshold scheme is generally used to sparsify the PC-based FBN by filtering out the noisy or weak connections. For detailed discussion on the thresholding strategy, please refer to Section 3.2.1 in [29].

### B. Partial Correlation

Partial correlation is used in FBN estimation for treating the confounding problem involved in the full correlation methods such as PC [30]. A general approach to calculate partial correlation is based on the estimation of inverse covariance matrix [31]. However, this approach may be ill-posed due to the singularity of the estimated sample covariance matrix **Σ = X^*T*^X**. To alleviate this issue, a regularization term *R*(**W**) is generally introduced into the FBN estimation model as follows.

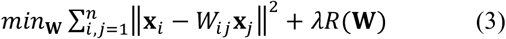

Equivalently, it can be further simplified to the following matrix form:

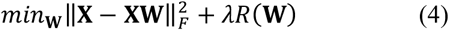

In Eq. (3) or (4), the first term implies to invert the covariance matrix Σ [31], and *λ* is a regularized parameter to control the balance between the first (data-fitting) term and the second (regularization) term. The most popular regularizer is *l*_1_-norm, i.e., 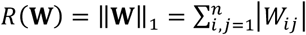, which corresponds to the sparsity prior of FBN, and leads to the LASSO or sparse representation (SR) model [15,32,33]. As discussed previously, some other regularizers have been investigated in recent years to encode different priors, which, however, is beyond our main focus of this paper.

### C. A Regularized FBN Estimation Framework

According to a recent study [12], a large family of FBN estimation models can be summarized by the following regularized framework:

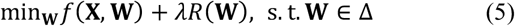

where *f*(**X, W**) is a data-fitting term, aiming to capture some statistics of the data (e.g., the covariance or inverse covariance structure), and *R*(**W**) is the regularization term, aiming to encode the biological/physical priors of the FBN. Sometimes, specific constraints (e.g., symmetry or positive semi-definite) are included in Δ for shrinking the search space of **W** towards better FBNs.

Such a regularized framework can *not only* stabilize the statistical estimation, *but also*, more importantly, provide a general platform for designing the new FBN estimation method in this paper.

## III. THE PROPOSED METHOD

As mentioned earlier, both prior information and data quality have a significant influence on the performance of the estimated FBNs. The priors can be generally encoded by the regularization terms. In contrast, however, the quality of time points in the fMRI series cannot be easily measured without the guidance of a specific learning/estimation task. Therefore, in this section, we propose a novel solution for identifying the quality of time points with the FBN estimation (task) together, by which we expect to get cleaner data, and, in turn, more accurate FBNs.

### A. Motivation

Before presenting the novel FBN estimation model, we first introduce the motivation and the basic idea behind it by a toy example. As shown in Fig. 1, we have a set of fMRI series from *N*=5 ROIs, each of which includes *T*=30 time points. According to the previous discussion, even a sophisticated preprocessing pipeline cannot guarantee all the *T* time points in the fMRI series are clean. To illustrate this more clearly, some data points in Fig. 1 are labelled as “clean”, while the others are labelled as “dirty”. Equivalently, a binary indicator can also be used to denote the quality of the data, with 0 corresponding to “dirty”, and 1 to “clean”.

**Fig. 1.**
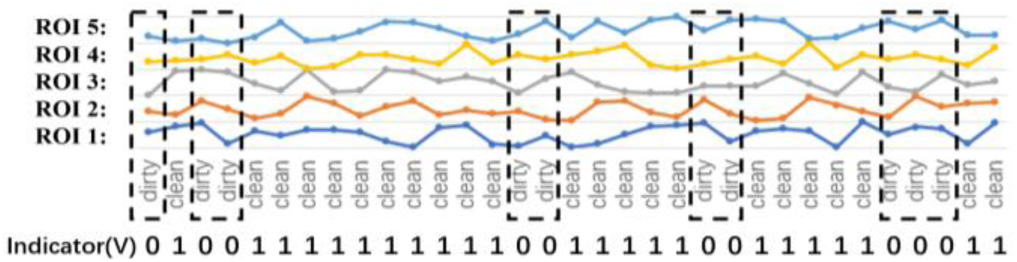
A toy example for illustrating the motivation of the proposed method.

However, the indicator cannot be observed directly in the real-world applications. Traditionally, a data-scrubbing operation is used to identify the quality of the data points (or, equivalently, determine the value of the indicator) by detecting the micro head motion [21]. Despite its potential effectiveness (also with some debates [23-25]), such a scheme 1) cannot remove all the “dirty” data points caused by different factors; 2) tends to scrub too many time points due to the lack of a control mechanism [12,21]; 3) is independent on the specific task, and thus will not necessarily benefit the ensuing FBN estimation.

### B. Model

To address the above problems, in this section, we propose a task-dependent data scrubbing scheme for FBN estimation. In particular, we consider the binary indicator as a variable *v*_*t*_for denoting the quality of the *t*th time point (*v*_*t*_ = 0 for “dirty”, and *v*_*t*_ = 1 for “clean”), and thus the regularized FBN estimation framework in Eq. (5) can be extended to the following form with an indicator variable.

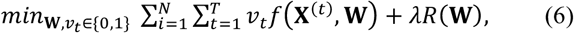

where **X**^(*t*)^ is the *t*th row of the data matrix **X**. Note that, when the indicator variable *v*_*t*_ = 0, the *t*th time point in the fMRI series will be removed, meaning that it has no contribution to the FBN estimation; when *v*_*t*_ = 1 for all *t* = 1, 2, …, *T*, Eq. (6) will reduce to the original FBN estimation framework given in Eq. (5).

However, in practice, Eq. (6) will always get a trivial solution (i.e., *v*_*t*_ = 0 for all *t* = 1, 2, …, ^*T*^), since it can really minimize the objective function. Therefore, we introduce a negative *l*_1_-norm regularization term, 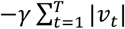, into Eq. (6). Additionally, we relax1 the binary indicator variable *v*_*t*_ϵ {0,1} to a real range [0,1], for simplifying the solving of the optimization problem. Then, we get the following model:

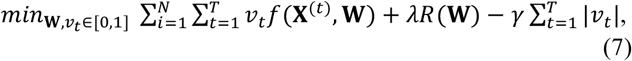

where *λ* and *γ* are two regularized parameters for controlling the balance among the three terms in the objective function. Especially for *γ*, it plays a role to determine the number of removed time points from the whole series. When *γ* has a small value close to 0, most of the time points will be removed. On the other hand, when *γ* has a large value, all the time points in the series will be kept for FBN estimation. In other words, the proposed model can scrub the data adaptively in the process of the FBN estimation, by controlling the hyper-parameter *γ* and learning the indicator variable *v*_*t*_from the data. Therefore, we name our proposed scheme as FBN estimation with self-scrubbing (**SS** model for short).

In principle, any data-fitting term can be used to realize the SS model, but, in this paper, we adopt the partial correlation strategy shown in Eq. (3) or (4), since it overcomes the confounding effect and is empirically verified to be more effective [11] than the full correlation. Consequently, the partial correlation based self-scrubbing FBN estimation model is given as follows:

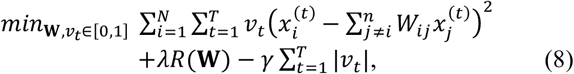

where *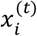* is the *t*th time point of the series associated with the *i*th node (ROI). For simplicity, we rewrite Eq. (8) into the following matrix form:

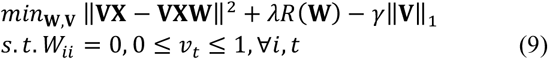

where V = diag(*v*_1_, *v*_2_, …, *v*_*T*_) ϵ *R*^*T×T*^ is a diagonal matrix containing the indicator variables on its principal diagonal. The constraint, *W*_*ii*_ = 0, is employed only to avoid the trivial solution that leads to **W** being an identity matrix. For *R*(**W**), we can, in principle, use any off-the-shelf regularizers, such as *l*_1_-norm, _*l2,1*_-norm [18,36], trace norm and their combination. However, this problem goes beyond our main focus in this paper. Therefore, we only attempt the *l*1-norm (sparsity prior) due to its simplicity and effectiveness [32], and get the specific FBN estimation model (named **SR+SS**) as follows.

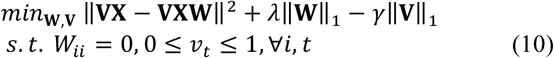

### C. Optimization Algorithm

Considering that there are two variables **V** and **W** involved in Eq. (10), in this paper we employ the alternative convex search (ACS) [37] method to solve them alternately by the two following steps. Before that, we first initialize the indicator **V** as an identity matrix, meaning that we select all time points in the first iteration.

**Step 1**: With a fixed **V**, Eq. (10) reduces to a traditional FBN estimation problem that can be solved by many convex optimization methods. Here, we use the proximal method [38] due to its efficiency and simplicity. Two main steps are involved in the proximal method, including gradient descent and proximal operation. First, for the data-fitting term 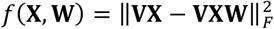 whose gradient w.r.t W is ∇_**W**_***f*(X, W)** = 2**X**^*T*^ **V**^*T*^**VXW** − **X**^*T*^ **V**^*T*^**VX**, we have the following update formula according to the gradient descent criterion:

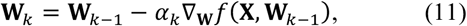

where *αk* denotes the step size of the gradient descent. Then, the proximal operator is imposed on the current **W**. For the sparsity regularizer *λ*∥W∥_1_, the proximal operator is defined [38] as follows.

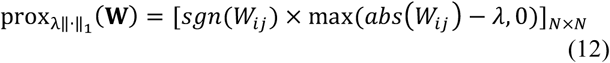

where *sgn*(*W*_*ij*_) and *abs*(*W*_*ij*_) return the sign and absolute value of *W*_*ij*_, respectively.

**Step 2**: With a fixed **W**, we update **V**. Now, Eq. (10) reduces to the following optimization problem,

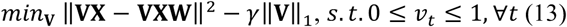

or

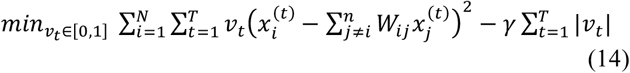

Eq. (14) can be further simplified to the following problem.

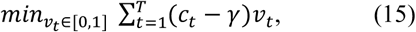

Where 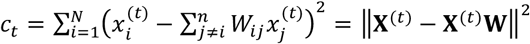 is a constant. We note that Eq. (15) is a linear programming problem, and thus can easily get its optimal solution as follows.

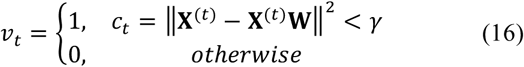

This means that 1) if ‖X^(*t*)^ - X^(*t*)^W^2^ > *γ*, the *t*th time point is more likely to be removed by labelling it with 0; on the contrary, 2) if ‖**X**^(*t*)^ - **X**^(*t*)^**W**^2^ < *γ*, the *t*th time point will be kept by labelling it with 1. Such a formula for updating *v*_*t*_coincides well with the intuition that the “dirty” time points (labelled as 0) can cause a poor fitting with high bias/residual (i.e., ‖**X**^(*t*)^ - **X**^(*t*)^**W**^2^ > *γ*). In fact, in the experimental section, we will further illustrate this problem based on a simulated dataset, and empirically verify that the proposed method can automatically detect the “dirty” time points. Finally, we summarize the algorithm for solving Eq. (10) in ALGORITHM I.

**ALGORITHM I.**
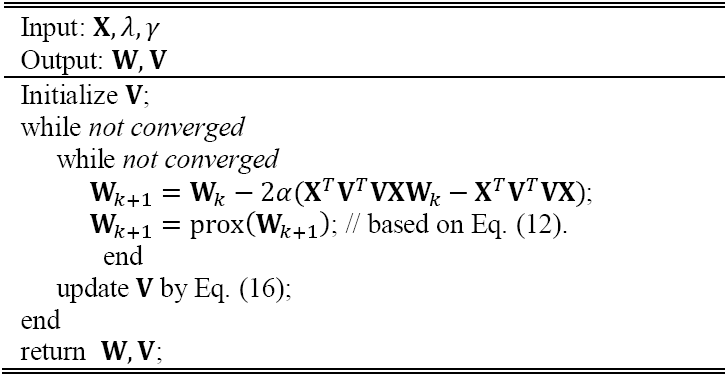
ESTIMATING FBN WITH SELF-SCRUBBING

## IV. EXPERIMENTS

In this section, we first illustrate how the proposed method works by an experiment on simulated data in Section <u>A</u>, and then verify its effectiveness by identifying Mild Cognitive Impairment (MCI) from Normal Control (NC) based on the estimated FBNs in Section <u>B</u>.

### A. An Illustrative Experiment on Simulated Data

For the convenience of interpretation and visualization, in this example, we only consider the simplest case that *N*=2. Thus, we have two time series, *x*_1_ and *x*_2_, associated with ROI_1_ and ROI_2_, respectively. Without loss of generality, we suppose that there is a strong connection between ROI_1_ and ROI_2_. For simulating the strong connection (i.e., correlation), we generate the data by an approximately linear relationship that *x*_1_≈ 5 × *x*_2_, as shown in Fig. 2(a) and (b). Based on the generated data, we can easily calculate the correlation coefficient between ROI_1_ and ROI_2_, and the result is 0.985.

**Fig. 2.**
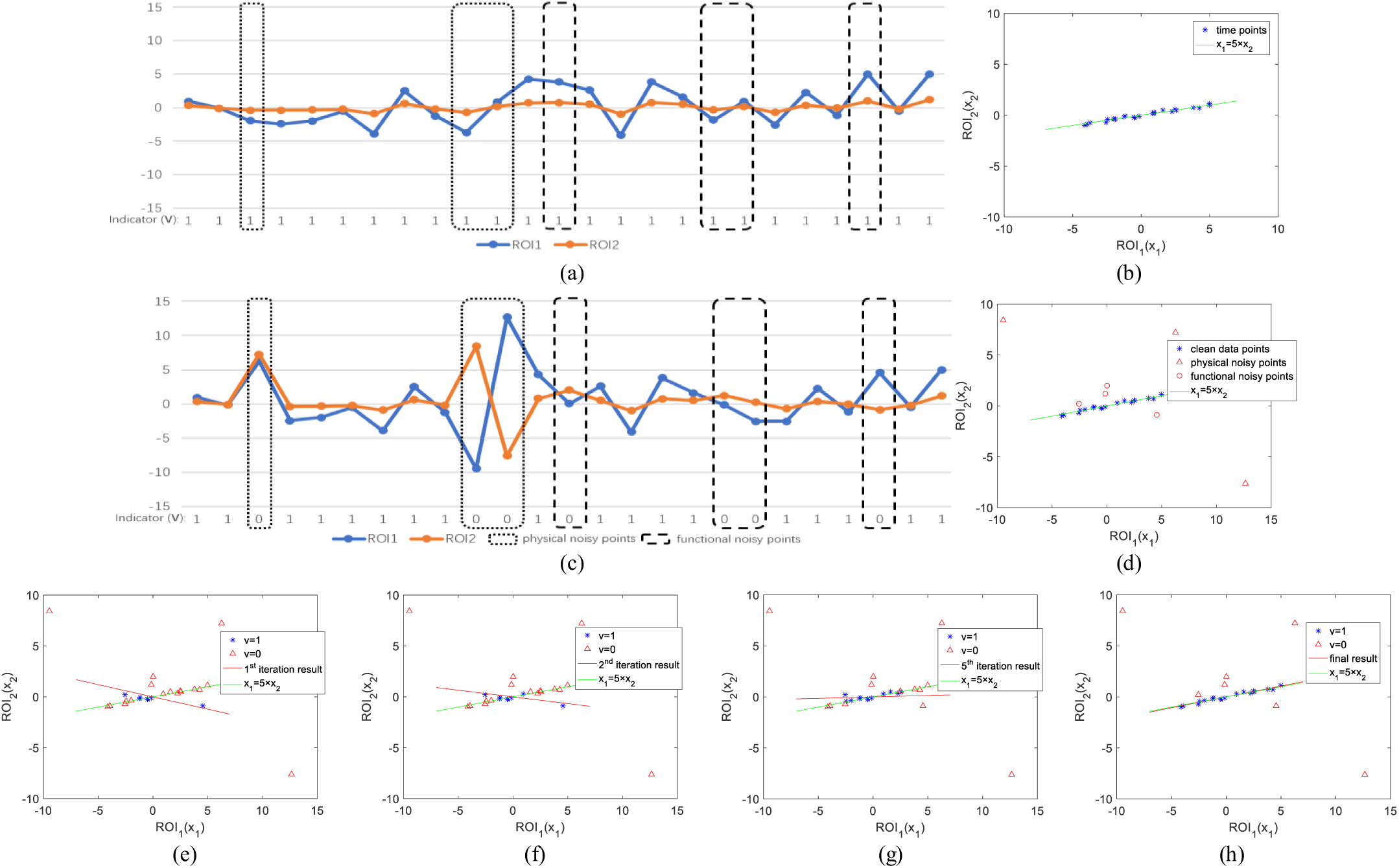
A toy example for illustrating how the proposed method works. (a) Original ideal signals simulated without noises; (b) scatter plot corresponding to (a); (c) observed signals simulated by introducing 7 noisy points into the original signals in (a); (d) scatter plot corresponding to (b); (e)-(h) the iterative results of the proposed algorithm. Note that the proposed method can effectively remove the noisy points gradually with the iterations.

However, as discussed previously, the correlation can be significantly affected by the data quality. To illustrate this, we change several time points randomly in the generated data for simulating the artifacts or noises. In particular, as shown in Fig. 2(c) and (d), we 1) introduce a large-amplitude shake in the first three “dirty” points to simulate the physical noises, since the physical noises (e.g., motion artifacts) have a high relation with the large-amplitude changes in signals [34,35]; 2) introduce four “dirty” points to simulate the possible functional noises, by setting them off the main direction (i.e., *x*_1_ = 5 × *x*_2_). Here, we simply consider the functional noises as non-resting-state signals that lie in the normal range, but deviate from the resting-state signals, according to a recent study [39].

Now, the correlation between ROI_1_ and ROI_2_ is -0.399, meaning that even limited change of data points can have a big influence on estimation of the functional connection. Based on the simulated data with “dirty” time points, we run the proposed algorithm, and find that it can remove the “dirty” points gradually with the iterations, as shown in Fig. 2(e) - (h).

### B. MCI Identification

#### 1) Data Acquisition and Preprocessing

In this study, we validate the proposed method by MCI and NC classification on two publicly available datasets. One is from Alzheimer’s Disease Neuroimaging Initiative (ADNI) ^2^, and the other is from Neuroimaging Informatics Tools and Resources Clearinghouse (NITRC)^3^ shared by a recent study [12].

For ADNI dataset, 110 participants, including 51 MCIs and 59 NCs, are adopted in this experiment. The fMRIs are obtained by 3.0T Philips scanners with the following parameters: TR/TE = 3000/30mm, flip angle = 80, imaging matrix=64×64, 48 slices, 140 volumes, and voxel thickness = 3.3mm. SPM8 toolbox ^4^ and DPARSFA (version 2.2) [40] are used to preprocess the fMRI data according to the well accepted pipeline. The first 10 R-fMRI volumes of each subject are discarded to avoid signal shaking. The remaining images are first corrected for different slice acquisition timing and head motion [41]. Then, regression of ventricular and WM signals as well as six head-motion profiles are conducted to further reduce the effects of nuisance signals. Mean fMRI series of each ROI is band-pass filtered (0.01-0.08Hz). Depending on the automated anatomical labeling (AAL) atlas [42], the pre-processed BOLD time series signals are partitioned into 116 ROIs. At last, we put these time series into a data matrix **X** ϵ *R*^137^×^116^.

For NITRC dataset, it includes 46 MCIs and 45 NCs that come from the participants recruited via advertisements in local newspapers and media. A similar preprocessing pipeline [12] are employed as in the ADNI dataset, except that a scrubbing operation is applied, for NITRC dataset, to remove the time points with frame-wise displacement larger than 0.5 [21]. We mainly use these two datasets for verifying that our proposed method can work well, whether the fMRI series are scrubbed or not.

#### 2) FBN estimation

After obtaining the preprocessed fMRI data, we estimate FBNs based on three different methods, PC, SR, and the proposed SR+SS. In Fig. 3, we visualize the adjacency matrices5 of the FBN estimated by these methods. For SR, we simply set the regularized parameter λ = 1, and for SR+SS, λ = 1 and γ= 0.5.

**Fig. 3.**
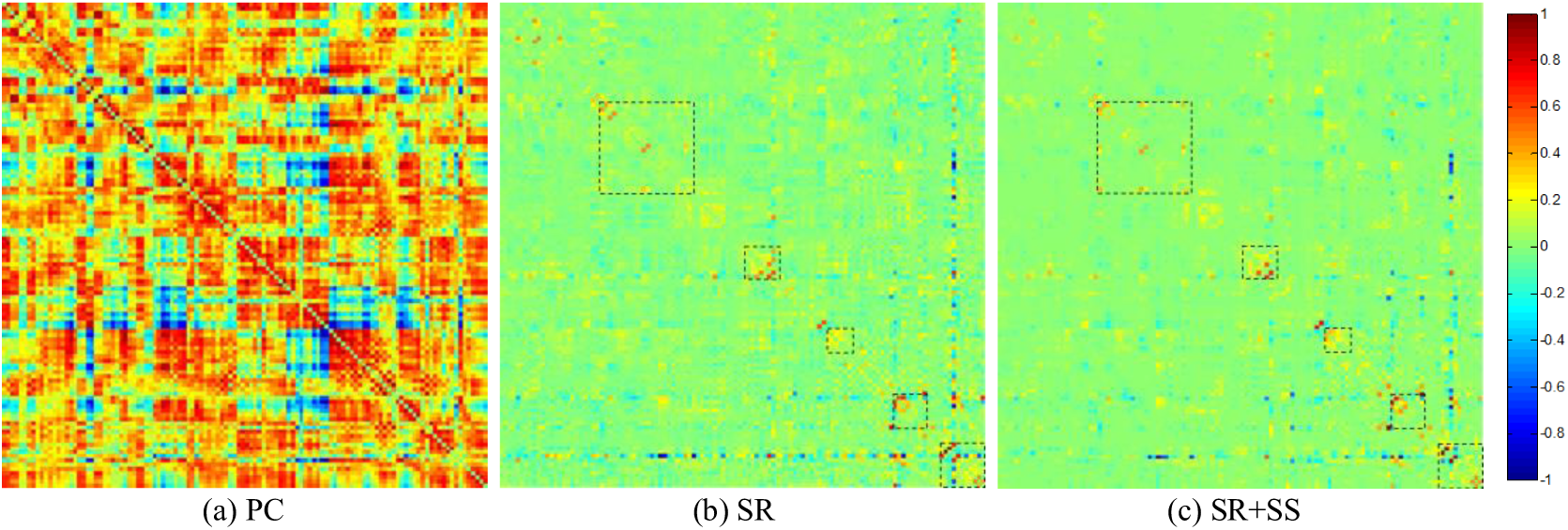
The FBN adjacency matrices of a subject estimated by 3 different methods. The rectangles in (b) and (c) help for distinguishing the differences between the two figures.

As shown in Fig. 3, the FBN estimated by PC has a topology highly different from that of the partial correlation-based methods (i.e., SR and SR+SS), since they use different data fitting term. In contrast, SR and SR+SS lead to a similar FBN structure by using the same kind of data fitting term. The differences of the FBNs estimated by SR and SR+SS methods lie mainly in several specific brain regions. In order to examine which brain regions have changed the connections following the self-scrubbing, we simply sum up the connection weights of all the subjects for each region. The result is shown in Fig. 4, where the height of each bar represents the sum of connection weights changing across different brain regions.

**Fig. 4.**
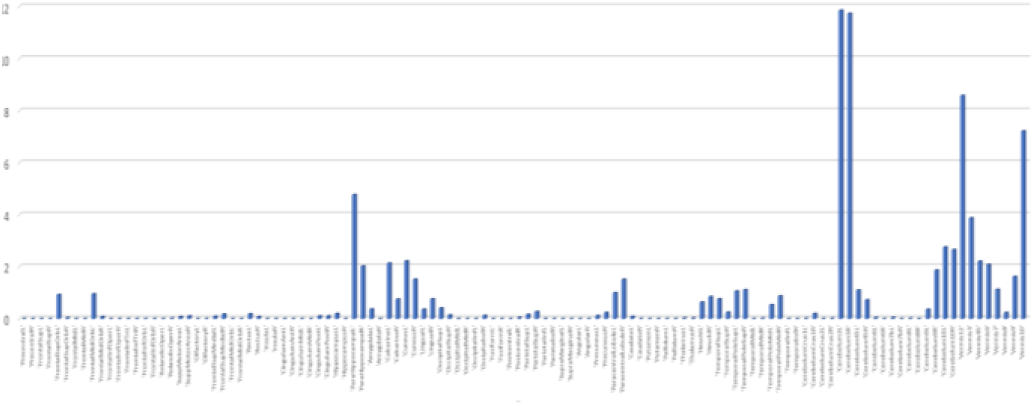
The connection weight change of FBN on each node (brain region) between SR and SR+SS. We first calculate the absolute value of the difference between the connections estimated by SR and SR+SS respectively. Then, we sum up the different weight values of the subjects for each brain region.

As we can see in Fig. 4, by scrubbing the “noisy” time points, a large number of changes happen in cerebellum regions, which is possibly related to the head motion or some other tiny movements. Also, the changes take place in hippocampus and frontal brain regions, which may be related to the psychological phenomena such as mind-wandering.

#### 3) Feature selection and Classification

Once we obtain the FBNs of all subjects, the subsequent task is to classify the MCI and NC based on the estimated FBNs. Then, the problem turns to determine which features and classifiers should be used for classification. Considering the big influence of different steps in the classification pipeline on the final accuracy [18], it is difficult to conclude whether the FBN estimation methods or the ensuing feature selection and classification methods contribute to the ultimate result. Therefore, we only adopt the simplest feature selection method (t-test with *p<*0.01) and the most popular SVM [43] classifier (linear SVM with default parameter *C=*1) in our experiment.

Due to limited samples, we test the involved methods using the leave-one-out cross validation (LOOCV) strategy, in which only one subject is left for testing while the others are used to train the models and get the optimal parameters. For the choice of the optimal parameters, an inner LOOCV is further conducted on the training data by a grid-search. For the regularized parameter *λ*, the candidate values range in [2^-5^, 2^-4^, …, 2^0^, …, 2^4^, 2^5^]; for the regularized parameter *γ*, the candidate values range in [0.1,0.2, …,0.9,1]; for the threshold in PC, we use 20 sparsity levels from [5%, 10%, …, 95%, 100%], where, for example, 90% means that 10% of the weak edges are filtered out from the FBN.

A set of quantitative measurements, including accuracy, sensitivity and specificity, are used to evaluate the classification performance of different methods. The classification results corresponding to these methods on ADNI and NITRC datasets are given in Fig. 5. Based on the results, we observe that the proposed SR+SS method achieves the best performance in accuracy, sensitivity and specificity on both ADNI and NITRC datasets. This illustrates that the self-scrubbing strategy can improve the performance of SR method in both scrubbed and non-scrubbed datasets.

**Fig. 5.**
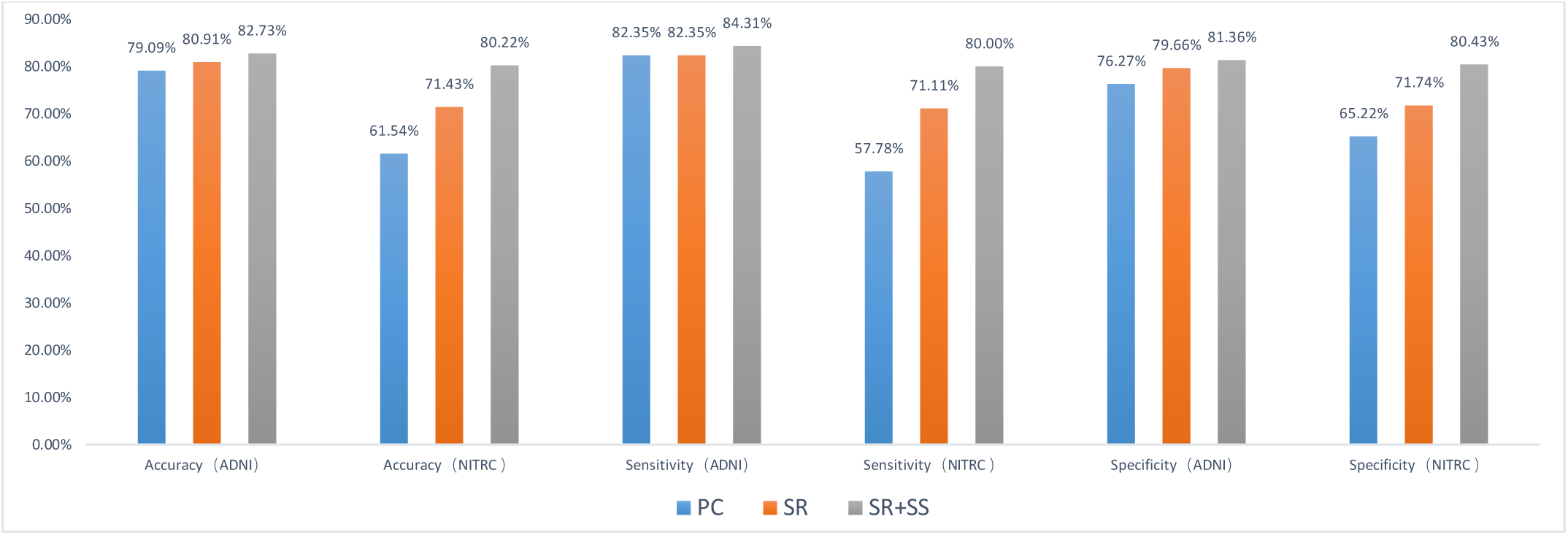
Comparison of classification results based on three different methods. The results obtained by LOOCV test show that the proposed SR+SS achieves the best performance.

## V. CONCLUSION

The observed fMRI time series commonly contain various artifacts or noises, thus leading to a poor estimation of the FBN. In this paper, we propose a novel FBN estimation method by incorporating a latent variable into the regularized FBN estimation model as an indicator of the data quality. Then, we design an alternating optimization algorithm to solve the new model. As a consequence, the proposed method can estimate FBN and scrub the fMRI series adaptively in a single framework. In particular, we adopt the SR as a simple test platform in this paper for developing our method that is then validated on simulated and real-world (both scrubbed and non-scrubbed) datasets. The experimental results show that our proposed method significantly outperforms the baseline methods. Also, in order to examine how FBN changes following the self-scrubbing, we compare the FBNs estimated by SR and SR+SS methods. The result shows that changes mainly happen in the high-order brain regions (e.g., hippocampus and frontal brain region) and cerebellum region, which illustrates that “dirty” points may be highly related to motion artifacts and functional processes.

The proposed method still has several shortcomings that need to be improved. For example, we select all time points in the first iteration, which may lead to a bad regression result (local minimum) when a large number of “dirty” points exists. In addition, the possibly useful information from the removed points can be lost, since the proposed method simply discards them based on the hard indicator. In the future, we plan to develop a soft (or probabilistic) version of the proposed method for higher flexibility. It is also worth pointing out that the proposed self-scrubbing strategy can be easily transferred into the other FBN estimation models with any combination of data-fitting and regularization terms. Therefore, we will try more combinations towards better accuracy in the future.

## Footnotes

1 In Section C, we will find that such a relaxation is tight, thus resulting in a binary solution for *v*_*t*_.

2 http://adni.loni.ucla.edu

3 http://www.nitrc.org/projects/modularbrain/

4 http://www.fil.ion.ucl.ac.uk.spm

5 The elements of the adjacency matrix indicate the connection strengths of the node pairs in the network. Here, for the convenience of comparison among different methods, all the weights are normalized to the interval [-1 1].

